# A hybrid mesh and voxel based Monte Carlo algorithm for accurate and efficient photon transport modeling in complex bio-tissues

**DOI:** 10.1101/2020.10.01.322982

**Authors:** Shijie Yan, Qianqian Fang

## Abstract

Over the past decade, an increasing body of evidence has suggested that threedimensional (3-D) Monte Carlo (MC) light transport simulations are affected by the inherent limitations and errors of voxel-based domain boundaries. In this work, we specifically address this challenge using a hybrid MC algorithm, namely split-voxel MC or SVMC, that combines both mesh and voxel domain information to greatly improve MC simulation accuracy while remaining highly flexible and efficient in parallel hardware, such as graphics processing units (GPU). We achieve this by applying a marching-cubes algorithm to a pre-segmented domain to extract and encode sub-voxel information of curved surfaces, which is then used to inform ray-tracing computation within boundary voxels. This preservation of curved boundaries in a voxel data structure demonstrates significantly improved accuracy in several benchmarks, including a human brain atlas. The accuracy of the SVMC algorithm is comparable to that of mesh-based MC (MMC), but runs 2x-6x faster and requires only a lightweight preprocessing step. The proposed algorithm has been implemented in our open-source software and is freely available at http://mcx.space.

## 1. Introduction

With the rapid emergence of powerful parallel computing platforms, especially those of graphics processing units (GPUs), the Monte Carlo method (MC) has been gaining popularity in light transport modeling within bio-tissues due to its high accuracy and scalability in computation [1,2]. As a stochastic solver to the radiative transfer equation, MC provides superior accuracy for general complex media, including low-albedo tissues where the diffusion approximation fails, such as cerebrospinal fluid (CSF) in the brain, lungs, and synovial fluid in the joints.

A range of media discretization strategies have been explored in MC simulations over the past two decades, driven by the desires to accommodate increasing complexities in imaging heterogeneous and multi-scaled tissues. As one of the most widely used MC algorithms, MCML was developed to simulate only multi-layered media [3]. The voxel-based MC (VMC) was proposed to handle arbitrarily heterogeneous media [4] by representing tissue domain using a three-dimensional (3-D) Cartesian grid of cubical voxels. VMC has seen hundredfold acceleration on GPUs [2, 5], largely owing to its inherent simplicity of the underlying data structure as well as low computational overhead in memory access. In recent years, several studies have pointed out that the orthogonal axis-aligned boundaries in a voxelated domain could produce unexpected modeling errors due to the unique optical characteristics associated with the special orientations of the boundary facets [6]. Recently, in [7], we have identified two types of artifacts that may arise when adopting a voxelized domain representation in MC. First, rasterization of a curved surface results in deviations between the original surface and a voxelated surface, as shown in Fig. 1(a) (referred to as the Type I error hereinafter). We found that this type of error can generally be reduced when a refined grid is used, shown in Figs. 1(b)–1(c), with a cost of dramatically increased computation. The second type of error (Type II) was caused by the distinct optical characteristics due to reflection/refraction of a voxelized surface compared to the smooth surface before discretization, as also highlighted in [8]. In such a case, progressively refining the grid fails to reduce the Type II error because this discrepancy is inherent to the surface shape.

**Fig. 1.**
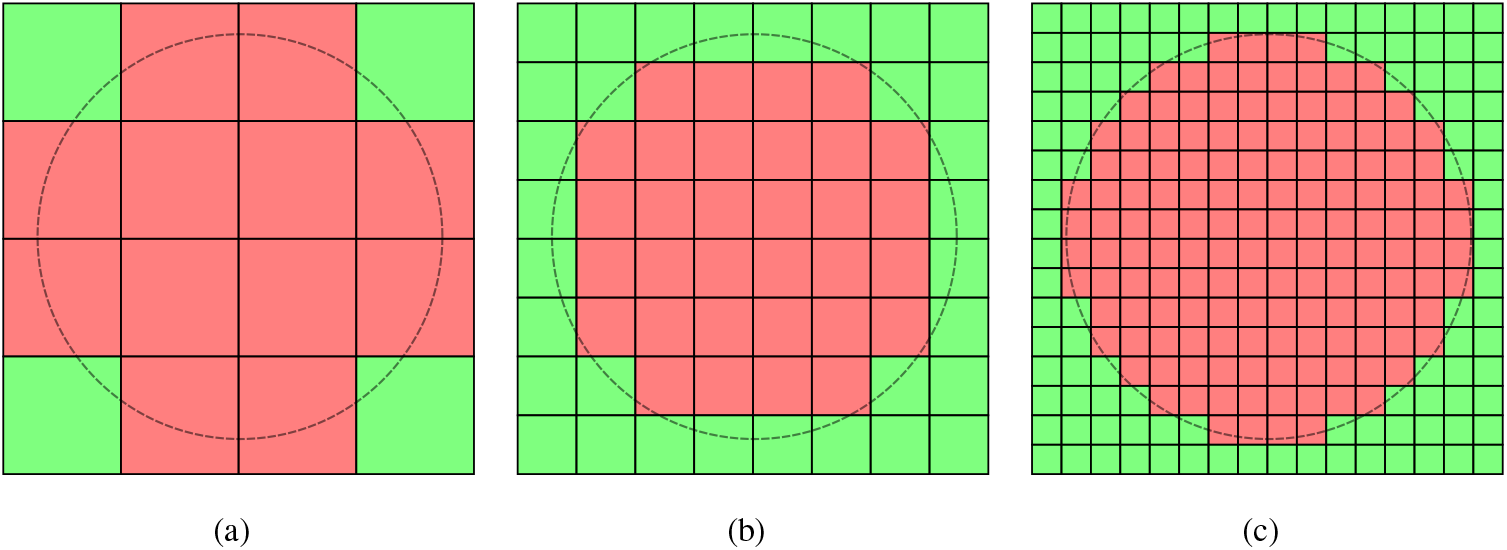
An illustration to show domain rasterization errors (Type I) of a curved interface (grey-dashed line) at three voxel sizes. The color shades represent rasterized tissue regions. Such mismatch can be reduced via voxel refinement, as shown in (b) and (c).

In the past years, considerable efforts have been invested to improve MC accuracy for handling complex tissue boundaries. A shape-based MC approach was proposed to improve modeling accuracy by considering parametrically defined 3-D shapes [8–11], which are largely limited to simple geometries such as spheres and cylinders [1]. A surface MC approach was explored in [12, 13], inspired partly by contemporary computer graphic rendering techniques. In this approach, triangular surfaces are used to separate the space into piece-wise-homogeneous tissue domains. A photon packet is cast as a ray and subsequently intersects with triangular patches, in a similar manner to ray-tracing in computer graphics. By further tessellating the domains between triangular surfaces, the mesh-based MC (MMC) using tetrahedral meshes was further proposed [14, 15] to dramatically accelerate the ray-triangle intersection testing. However, creating anatomically accurate tetrahedral mesh is not a trivial task and often requires many careful considerations and dedicated meshing tools [7]. Furthermore, a mesh based data structure is significantly more complex and memory-expensive when compared to VMC. This can potentially result in reduced efficiency when implemented on memory-sensitive hardware such as GPUs [16, 17]. To take advantage of the low overhead of the voxelated data structure, a refined VMC approach was recently proposed [18] to incorporate surface normal information, pre-computed using a gradient operator, at the boundary voxels. Although this method has showed improved accuracy in handling boundary reflections compared to voxelated boundaries, it does not address the aforementioned Type I error caused by the partial-volume effect between boundaries, especially those between refractive-index matched tissues.

In this work, we describe a hybrid mesh and voxel MC algorithm, referred to as the “split-voxel” MC or SVMC hereinafter, that is capable of accurately modeling light transport across curved boundaries while utilizing the highly efficient voxelated data structure for ray-tracing computation. This hybrid surface-voxel representation is efficiently generated using a pre-processing step applicable to any existing voxelated domain by performing a fast marching-cubes [19] mesh generation process to create voxel-bounded triangular boundary patches. Surface normal information and partial-volume fractions of the boundary voxels can be subsequently derived from the marching-cubes surfaces within the boundary voxels. By packing the additional normal and oblique surface information in an extended voxel data format, we then further utilize this refined boundary information by revising the ray-tracing computation within a voxel to explicitly consider the oblique boundary interface if present. Non-boundary-intersecting voxels are handled the same way as in a VMC simulation, making SVMC nearly as accurate as MMC, yet without the overhead of tetrahedral mesh generation and accessing complex data structures.

In the remainder of this manuscript, we first detail the pre-processing steps to create voxel-bounded triangular meshes from an arbitrary input volume and our approaches to compute and encode the oblique boundary information within boundary voxels. We then describe an extended ray-tracing method to handle photon propagation within a boundary voxel using both voxel and the embedded oblique surface information. Further considerations for efficient implementations on our GPU-accelerated simulator, MC eXtreme (MCX) [5] are discussed, followed by the description of the input data layout and throughput optimization strategies. Next we validate the proposed algorithm and quantitatively demonstrate the significant reduction of both Type I and Type II errors in several benchmarks of heterogeneous domains. Finally, we discuss the limitations of this work and summarize our main findings.

## 2. Methods

In VMC, a heterogeneous tissue domain is typically represented by a 3-D multilabeled volume [4, 5], where a single integer is assigned to each voxel to represent the index of the tissue type located in that voxel. This voxelated representation is typically resulted from a segmentation process where a curved boundary is replaced by a terraced voxel surface despite that real-world tissue boundaries rarely align with orthogonal Cartesian planes. Such segmented data inherently contains discretization error that could potentially impact photon modeling accuracy. In order to improve the boundary accuracy in VMC, one must first restore the missing curved boundary information.

There are several known approaches to restore curved boundary information using a voxelated data array. In one approach, one can apply a surface extraction algorithm such as *ϵ*-sampling, as used in our brain mesh generation pipeline [7]. However, these extracted surface triangles are not bounded by the containing voxel. In order to attribute the surface information to each intersecting voxel, one must perform a series of “surface Boolean” operations by “slicing” the surface with the bounding box of each voxel. While this is doable, the associated computational overhead can be quite high.

On the other hand, the marching-cubes mesh generation algorithm [19] is known to be highly efficient when reconstructing curved surfaces from a 3-D volume. Aside from being fast, the marching-cubes algorithm outputs triangles with boundaries naturally aligned with the bounding box of the enclosing voxel, making it highly convenient and efficient to further process. We want to point out that the marching-cubes surface triangles extracted directly from a binary or multi-labeled volume may only present limited orientations, such as 45° or 90°. Although they can be oblique, the limited orientation of the triangle patches remains a source of discretization error. To create a smooth boundary, a 3-D Gaussian filter can be applied to the binary or multi-labeled volume before performing a gray-scale marching-cubes surface extraction. With this pre-smoothing step, the qualities of the marching-cubes surface triangles are greatly improved, resulting in smooth and continuous orientation variations.

In Figs. 2(a-c), we compare different shape representation strategies using a spherical domain as an example. Compared to the voxelated surface in Fig. 2(a), the marching-cubes surfaces [Figs. 2(b-c)] show improved smoothness, especially when a 3-D Gaussian filter is used. In Figs. 2(d,e), we show magnified views of the marching-cubes surfaces in (b) and (c) respectively, centered at a selected voxel (black cube). The limited surface orientation is demonstrated in Fig. 2(d) when the marching-cubes extraction is applied directly to a rasterized binary (0-1) spherical domain. In comparison, the curved shape of the sphere is well recovered in Fig. 2(e) when applying a Gaussian smoothing before running the marching-cubes, achieved by calling the `isosurface` function in MATLAB (Mathworks, Natick, MA).

**Fig. 2.**
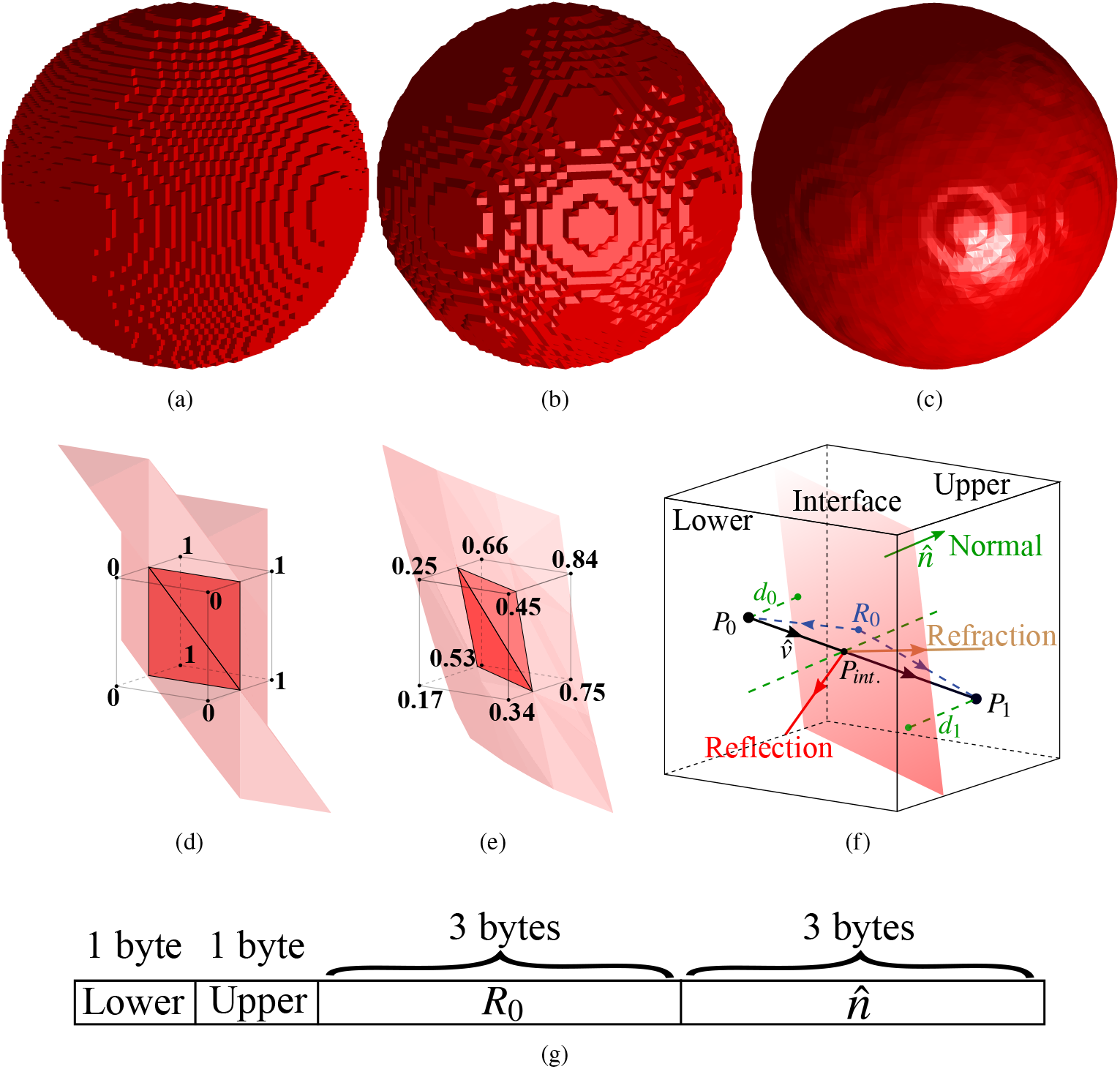
Illustrations of boundaries of a spherical domain extracted using (a) voxel representation, and marching-cubes (b) without and (c) with smoothing by a Gaussian-filter. Zoom-in views of the surfaces in (b) and (c) can be found in (d) and (e), respectively. We also show diagrams explaining (f) ray-tracing computation and (g) our extended voxel data memory layout that encodes additional shape information.

Note here that in order to facilitate the implementation and improve computational efficiency, two simplifications are applied in processing the marching-cube surfaces. First, marching-cubes can produce 2 to 4 disconnected surface components/patches [19] if the voxelated image was previously rasterized from scans of sub-voxel features, i.e. a shape feature that is smaller than the span of the voxel. In our implementation, we only extract the largest single-connected surface component and ignore other disconnected triangles. If this simplification should become a concern, one should consider increasing the resolution of the input volume in order to capture those fine details. Secondly, for each connected surface component, marching-cubes may produce up to 4 connected triangular patches [19]. To avoid storing large amount of mesh data in each voxel, here we approximate the marching-cubes per-voxel triangular surface by a single plane defined by a normal direction 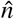, obtained using the area-weighted average from each connected triangles, and a single position *R*_0_ on the plane, computed as the centroid of the surface patch nodes, as illustrated in Fig. 2(f).

To store the additional shape information (*R*_0_ and 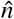), in Fig. 2(g), we show an extended voxel data format. In this new format, each voxel contains an 8-byte record (counting from the most to the least significant bits): byte 1 stores the tissue label (up to 256 types) of the current voxel (if not at a boundary), or, if at a tissue boundary, the tissue type at 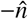 direction (i.e. lower); byte 2 stores the tissue type at 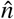 direction (upper) of the surface; bytes 3-5 store the 0-255 quantized *x/y/z* coordinates (only the decimal parts) of *R*_0_; bytes 6-8 store the 0-255 quantized *x/y/z* components of 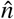 (each one between −1 and 1). This data format is optimized for GPU implementation to utilize the cache-line to accelerate reading from the global memory [20].

Subsequently, we develop an extended ray-interface intersection test to consider the intra-voxel boundaries, if present, as depicted in Fig. 2(f). In a non-boundary voxel, a photon packet propagates exactly the same way as in VMC. However, then it first enters a boundary voxel, indicated by a non-zero upper label (byte-2), at position *P*_0_, a dot product 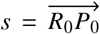 is performed to decide if the photon is in the “lower” (5 < 0) or “upper” (5 > 0) space of the interface. In the latter case, we set 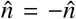 so that 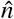 is always pointing to the region that is not “current”. Similar to the conventional VMC, photon path length *L*, is computed as the remaining scattering length or distance from *P*_0_ to voxel boundary, whichever is smaller [2,5]. For every step when a photon moves from 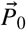 to 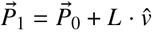, where 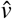 denotes the movement direction, we compute the signed distances from 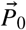 and 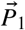, respectively, to the interface as 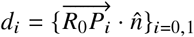, 1. When the photon path 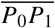 intersects with the interface, the inner product 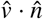 must be positive while *d*_0_ and *d*_1_ should have different signs, i.e. *d*_0_*d*_1_ ≤ 0. The photon is then moved to the position of the intersection point, 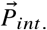, computed by

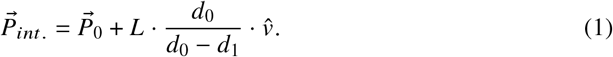

For refractive-index mismatched boundaries, the calculation of reflection and refraction is performed to update the propagation direction, similarly to MMC [14, 15]. Every time a photon transmits through the interface, we set 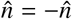 before the next intersection test, for the reason mentioned above.

## 3. Results and Discussions

In this section, we validate the proposed SVMC algorithm and quantify the accuracy improvement by comparing its solution with our GPU-accelerated mesh and voxel-based MC simulators: DMMC (dual-grid MMC) [15,17,21] and MCX (VMC mode) [5] in three heterogeneous domains. We first focus on validating the accuracy in two simple geometries. Benchmark B1 (“sphshells”) was previously described in Ref. [21]. Briefly, it contains a 60 × 60 × 60 mm^3^ cubic domain surrounded by air. Three concentric spheres, centered at (30.5,30.5,30.5) mm, with radii 10, 23 and 25 mm, respectively, divide the cubic domain into 4 layers. Different optical properties are assigned in each layer. From innermost to outermost, we have {*μ_a_* = 0.05/mm, *μ_s_* = 0.0/mm, *g* = 1.0 *n* = 1.37}, {*μ_a_* = 0.02/mm, *μ_s_* = 9.0/mm, *g* = 0.89, *n* = 1.37}, {*μ_a_* = 0.004/mm, *μ_s_* = 0.009/mm, *g* = 0.89, *n* = 1.37} and {*μ_a_* = 0.02/mm, *μ_s_* = 7.0/mm, *g* = 0.89, *n* = 1.37}, where *μ_a_* denotes the absorption coefficient, *μ_s_* denotes the scattering coefficient, g denotes anisotropy and n is the refractive index. We use this example to highlight the Type I error of VMC. Note that except for the air-tissue boundaries along the cube surface, all interior tissue boundaries are refractive-index matched. In benchmark B2 (“cubesph”), a spherical inclusion of radius 25 mm with *μ_a_* = 0.005/mm, *μ_s_* = 1/mm, *g* = 0.89 and *n* = 1.37 is embedded inside a 60 × 60 × 60 mm^3^ cube that is filled with air. We use the B2 benchmark to highlight the Type II error of VMC. In both B1 and B2, a pencil beam source is positioned at (30.5,30.5,0) mm pointing to the +*z* direction. In addition, to demonstrate the importance of performing the input volume smoothing prior to surface extraction, for all SVMC simulations, we test both hybrid domains derived with and without Gaussian smoothing. Next we expand our test to more realistic complex domains. The benchmark B3 (“brain19.5”) uses a brain segmentation of an MRI brain atlas (see Fig. 10(a) in Ref. [7]) and was used to illustrate both Type I and II errors in [7]. Additionally, we report several key optical parameters derived from the simulations and compare those with DMMC [21] and MCX reported previously (see Table 1 in Ref. [7]). For all benchmarks, 10^8^ photons are simulated on a desktop running Ubuntu 16.04 with an NVIDIA RTX 2080 GPU, and a Henyey-Greenstein phase function [3] is assumed. We compute the volumetric light fluence distributions using an output grid with 1 × 1 × 1 mm^3^ voxel resolution and report the simulation speeds (photons/ms) to characterize the impact on performance. All simulation scripts and input settings are available in our software repository (https://github.com/fangq/mcx) for reproducibility.

**Table 1.** Summary of simulated optical parameters, including the average photon partial pathlengths in the brain region (PPL_B_), total-pathlengths (TPL), and their percentage ratios (R_B_) derived from SVMC simulation of B3 (“brain19.5”). We also include the R_B_ reported by DMMC (reference) and MCX in [7].

| Det.<br># | SVMC |  |  | DMMC | VMC |
| --- | --- | --- | --- | --- | --- |
|  | PPL<sub>B</sub><br>(mm) | TPL<br>(mm) | R<sub>B</sub><br>(%) | R<sub>B</sub><br>(%) | R<sub>B</sub><br>(%) |
| 1 | 0.05 | 34.4 | 0.14 | 0.14 | 0.17 |
| 2 | 1.56 | 93.24 | 1.67 | 1.66 | 2.03 |
| 3 | 4.41 | 122.9 | 3.58 | 3.54 | 4.31 |
| 4 | 8.61 | 150.2 | 5.73 | 5.82 | 6.48 |
| 5 | 13.54 | 173.2 | 7.82 | 7.56 | 8.44 |
| 6 | 0.04 | 35.83 | 0.11 | 0.11 | 0.14 |
| 7 | 1.14 | 95.21 | 1.20 | 1.26 | 1.42 |
| 8 | 3.23 | 124.9 | 2.58 | 2.78 | 2.92 |
| 9 | 7.42 | 157.6 | 4.71 | 4.95 | 5.00 |
| 10 | 11.37 | 179.9 | 6.32 | 6.78 | 6.68 |

In Fig. 3, we show the cross-sectional contour plots to compare light fluence distributions between three MC algorithms for all benchmarks. For both B1 and B2 benchmarks [Figs. 3(a-b)], the fluence contour lines computed using SVMC with (red-dashed) and without (orange-dotted) Gaussian smoothing show improved accuracy over the conventional VMC (white-dashed). Between the two SVMC results, the one computed using smoothed volume shows significantly better match with the reference solution computed using DMMC (black-solid). In B3 [Fig. 3(c)], an excellent agreement between SVMC and DMMC solutions is observed along the scalp-air surface, suggesting that the Type II error was largely removed. However, the discrepancies in the CSF layer under detectors 7-9, a result of Type I error, was reduced but not eliminated. In Table 1, we further compare SVMC with VMC, using DMMC as a reference. Overall, metrics derived from SVMC are very similar to those from reference solutions (from DMMC) at all detectors, compared with those derived from VMC. Among all detectors, deviations between SVMC and DMMC appear to be smaller at detectors near the source – #1 to #3 and #6 to #7 – but becomes more noticeable at larger source-detector separations. We believe this deviation is a combined result of (1) the differences in mesh-generation algorithms (marching-cubes vs. *ϵ*-sampling used in Brain2mesh [7]), and (2) the surface approximations within a voxel in the SVMC.

**Fig. 3.**
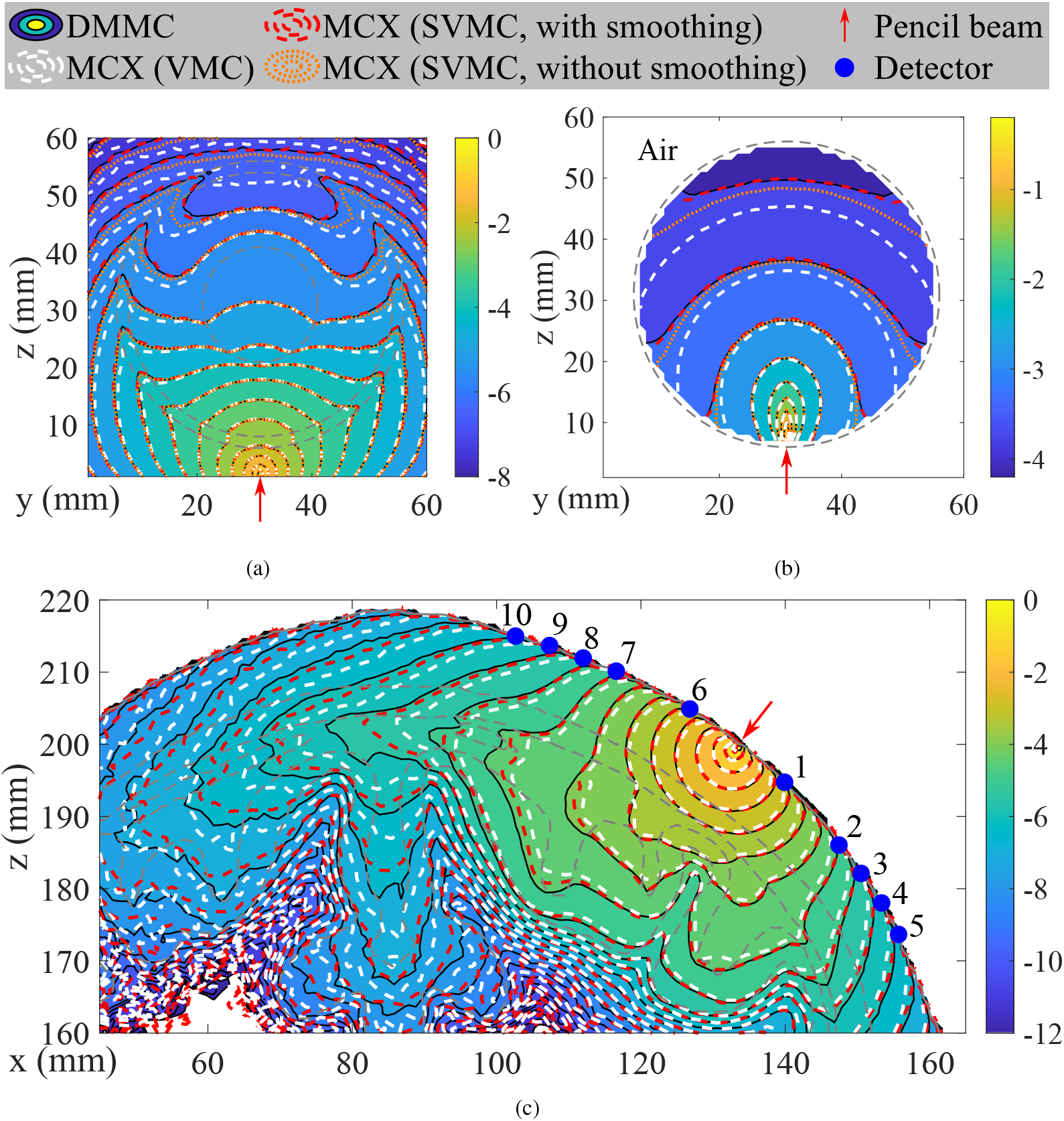
Fluence (mm^-2^, in log-10 scale) contour plots of dual-grid MMC (DMMC), MCX (conventional VMC mode) and MCX (SVMC mode with and without volume smoothing) in a range of benchmarks: (a) B1, (b) B2 and (c) B3. The red arrow represents an inward-pointing pencil beam source. Grey-dashed lines mark media boundaries.

In Table 2, the speeds of MCX in VMC and SVMC modes are both faster than DMMC in all benchmarks. Compared with the conventional VMC, SVMC reports 50.1%, 57.3% and 55.4% speed reduction, respectively, in benchmarks B1, B2 and B3 as a result of extra computation needed to consider the intra-voxel boundaries, along with the increased memory overhead due to the increased input data size. To quantify accuracy changes, we also compare the total absorption percentage – a ratio between the total absorbed and simulated energy. In B2 and B3 where tissue-air (mismatched) boundaries are curved surfaces, SVMC and DMMC report comparable results while VMC’s solution shows significant deviations. In addition, we compare the input data pre-processing time of SVMC and DMMC, with SVMC reporting 0.53 s, 0.26 s and 38.2 s and DMMC reporting 2.19 s, 1.39 s and 43.11 s for B1, B2 and B3 respectively. The pre-processing time of SVMC is largely associated with marching-cubes surface extraction and the computation of the interface geometric parameters. However, the marching-cubes algorithm is readily to be accelerated using GPU, potentially leading to significantly improved pre-processing speed for SVMC.

**Table 2.**
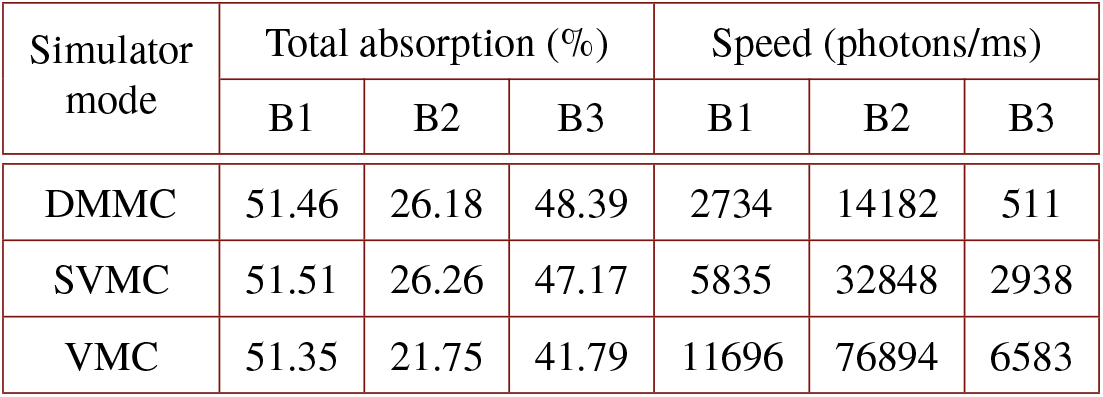
Summary of the total absorption and simulation speeds of DMMC (reference) and MCX (SVMC and VMC modes) for the selected benchmarks.

| Simulator mode | Total absorption (%) |  |  | Speed (photons/ms) |  |  |
| --- | --- | --- | --- | --- | --- | --- |
|  | B1 | B2 | B3 | B1 | B2 | B3 |
| DMMC | 51.46 | 26.18 | 48.39 | 2734 | 14182 | 511 |
| SVMC | 51.51 | 26.26 | 47.17 | 5835 | 32848 | 2938 |
| VMC | 51.35 | 21.75 | 41.79 | 11696 | 76894 | 6583 |

## 4. Conclusion

In summary, we present a hybrid MC simulation algorithm and data structure to improve voxel-based Monte Carlo photon transport by enhancing its capability to properly handle curved interfaces in 3-D complex media. Enabled by the marching-cubes algorithm, the surface mesh extracted from the volume can be efficiently processed and create hybrid data structure encoding intra-voxel boundaries. In addition, we have developed a fast ray-interface intersection testing algorithm and incorporated it into our GPU-accelerated MC simulator. Compared to conventional VMC, SVMC reports about 50% speed loss, but in return, it largely removed both Type I and II errors that are inherent to VMC. On the other hand, SVMC exhibits a level of accuracy nearly as good as MMC, but is 2x-6x faster and only requires a lightweight preprocessing. The SVMC algorithm has been implemented in our open-source MC simulator and is freely accessible at http://mcx.space.

## 5. Acknowledgment

This work is supported by the National Institute of Health under grants R01-GM114365, R01-EB026998, and R01-CA204443.

## 6. Disclosures

The authors declare no conflicts of interest.

## Notes

http://mcx.space

### Competing Interest Statement

The authors have declared no competing interest.

